# Investigating the role of the hippocampus in the implicit and explicit expression of statistical learning in patients with temporal lobe epilepsy

**DOI:** 10.64898/2026.09.04.749513

**Authors:** Emily Cordeiro, Daniela Herrera-Chaves, Nima Talaei, Iván Castro, Brent Hayman-Abello, Susan Hayman-Abello, Tara McAuley, Ana Suller-Marti, Stefan Köhler, Laura Batterink

## Abstract

Statistical learning (SL) has been proposed to depend on the hippocampus, but traditional neuropsychological theories of long-term memory posit that the hippocampus is only necessary for explicit memory processes, not implicit memory processes. To reconcile these two accounts, we exposed 27 temporal lobe epilepsy (TLE) patients to a continuous speech stream containing embedded trisyllabic words, then tested their statistical knowledge using both implicit and explicit behavioural measures. TLE patients showed impairments on two explicit measures of SL alongside intact performance on an implicit measure of SL. These results suggest that the hippocampus may only be necessary for the explicit expression, but not the initial acquisition or implicit expression, of statistical knowledge.

## 1 Introduction

### 1.1 Statistical learning

*Statistical learning* (SL) refers to the process of detecting recurrent patterns in the environment over time (Saffran, Newport, et al., 1996). By detecting these regularities, individuals can form predictions about future events, reducing cognitive demands by anticipating what is likely to occur next and allocating attention toward novel or unexpected information (Sherman et al., 2020). For example, orchestral pieces often develop through a sequence of musical passages in which instruments are introduced in a particular order, such as woodwinds followed by strings and then brass. After listening to the same piece many times, audience members may anticipate the moment when a particular instrument will begin playing while quickly noticing when a musician misses their entrance. Because these regularities are typically acquired incidentally, SL is regarded as an implicit learning process that occurs without conscious awareness (Perruchet & Pacton, 2006; Xu et al., 2023).

SL was originally studied in the context of language acquisition, where it was proposed to be a fundamental mechanism for speech segmentation. Spoken language is continuous and lacks consistently reliable acoustic markers (e.g., pauses, intonations) that indicate where one word ends and the next word begins, requiring infants and second-language learners to rely on some other cue to identify individual words. Alternatively, they may rely on *transitional probabilities*, or the statistical regularities between adjacent syllables. Specifically, syllables within words are more likely to occur together than syllables between words across a given language inventory (Aslin et al., 1998). For example, in the phrase *pretty*+*baby*, the syllables *pre* and *tty* are more likely to occur together in the English language than *tty* and *ba*. In their seminal study, Saffran, Aslin, and Newport (1996) were the first to propose that learners segment speech by becoming sensitive to transitional probabilities.

To test this hypothesis, they exposed 8-month-old infants to a two-minute artificial speech stream containing four trisyllabic nonsense words that repeated continuously in a pseudorandom order, such that the only cue to word boundaries was a reduction in the transitional probability between syllables of neighbouring words (e.g., *bidaku*+*padoti*+*golabu*+*bidaku*…). Following exposure, the infants were presented with two types of trials: repetitions of a word that was presented in the stream (e.g., *bidaku*) or repetitions of a foil item composed of three syllables that appeared together in the stream but crossed a word boundary (e.g., *ku*+*pado*). Looking times indicated that infants could distinguish between the two item types, showing a novelty preference for foil items. This provided the first behavioural evidence that statistical regularities alone were sufficient to provide word-boundary cues.

Since then, research over the last 30 years has demonstrated that SL is not limited to auditory speech segmentation but instead operates across many domains. Sensitivity to statistical regularities has been observed across other sensory modalities, including vision (Fiser & Aslin, 2001; 2002), touch (Conway & Christiansen, 2005) and cross-modal contexts (Conway & Christiansen, 2006; Mitchel & Weiss, 2011), as well as with nonlinguistic auditory stimuli (Saffran et al., 1999). Statistical regularities can also be extracted from more complex temporal structures, such as non-adjacent dependencies (Gómez, 2002), and from spatial configurations of visual stimuli (Fiser & Aslin, 2001). This ability has been demonstrated across the human lifespan, from infancy (e.g., Saffran, Aslin, et al., 1996; Teinonen et al., 2009), through childhood (e.g., Saffran et al., 1997; Arciuli & Simpson, 2011; Moreau et al., 2022), early adulthood (e.g., Batterink & Paller, 2017; Saffran et al., 1997; Wang, Köhler, et al., 2023), and into healthy aging (e.g., Wang, Köhler, et al., 2023). Consequently, SL is regarded as a fundamental mechanism that supports a wide range of perceptual and cognitive functions, including language acquisition, reading development, attention, object perception, and predictive processing (Bogaerts et al., 2020; Sherman et al., 2020).

### 1.2 Behavioural assessments of SL

Many studies investigating temporal SL continue to use paradigms adapted from the original artificial language paradigm developed by Saffran, Aslin, et al. (1996), whether they use auditory or visual sequences. These paradigms typically begin with an exposure phase during which participants passively listen to or view a continuous stream of stimuli. In the auditory-linguistic domain, streams typically consist of syllables grouped into pairs, triplets, or larger sequences – hereon referred to as “words” – that repeat in a pseudorandom order. Importantly, word boundaries are not obviously marked by pauses or other perceptual cues; instead, learners must rely primarily on changes in transitional probabilities between elements to identify the underlying structure of the stream. Following exposure, learning of the stream’s statistical regularities can be assessed behaviourally using a variety of memory-based tasks, which differ in the extent to which they rely on conscious retrieval of learned representations.

#### 1.2.1 Explicit expression of SL

Explicit memory refers to the conscious, intentional retrieval of previously encountered information, requiring intentional recollection or deliberate memory-based judgements (Tulving, 1985; Schacter, 1987; Schacter et al., 1989; Squire & Zola, 1996). Traditionally, the most widely used behavioural measure of SL has been the forced-choice recognition task (e.g., Saffran, Aslin, et al., 1996; Saffran, Newport, et al., 1996; Saffran et al., 1999; Fiser & Aslin, 2001; see Isbilen & Christiansen, 2022, for a meta-analysis), which is considered an explicit memory measure. In this task, participants are presented with a word that appeared during the exposure phase alongside one or more reconfigured foil items. Participants are asked to select the sequence that seems more familiar to them. Performance above chance indicates sensitivity to the statistical regularities embedded within the stream.

Another explicit measure of SL is the familiarity rating task (Batterink et al., 2015; Batterink & Paller, 2017). In this task, participants are presented with individual items – either a word that appeared during exposure or a reconfigured foil item – and they must rate how familiar each item seems to them using a Likert scale. If participants learned the statistical structure of the exposure stream, then they should rate words as being more familiar than the foil items.

Importantly, recent research suggests that knowledge acquired through SL can be expressed through both explicit and implicit forms of memory retrieval. Therefore, explicit tasks requiring intentional deliberation may not fully capture all aspects of statistical knowledge (Arcuili, 2017; Batterink et al., 2015; Bertels et al., 2012). To address this issue, many studies have also employed implicit measures of SL.

#### 1.2.2 Implicit expression of SL

In contrast to explicit measures, implicit measures of memory assess learning through changes in behaviour that result from prior experience without requiring conscious, intentional recollection of the learned information (Schacter, 1987, Schacter et al., 1989; Squire & Zola, 1996). In the context of SL, implicit measures typically assess sensitivity to the predictive structure of a learned sequence rather than requiring participants to make judgements about an encountered item. More recent studies of SL have implemented the target detection task, a reaction-time based measure of learning (Batterink et al., 2015; Turk-Browne et al., 2005). In this task, participants are presented with shorter segments of the exposure stream and instructed to detect a specific stimulus or “target” as quickly as possible via button press. Learning is indexed by faster responses to items that occur later within a word compared to initial items, as earlier items provide predictive information about upcoming items when the underlying statistical structure has been learned.

Evidence increasingly suggests that implicit and explicit expressions of SL reflect partially distinct forms of acquired knowledge. These measures show relatively low correlations with one another, follow different consolidation trajectories, and are differentially impacted by repeated testing, healthy aging, attention during learning, and neurological conditions (Batterink et al., 2015; Liu et al., 2023; Reyes et al., 2026; Wang, Köhler, et al., 2023; Wang, Rosenbaum et al., 2023). Therefore, understanding how these distinct behavioural expressions of SL relate to the different memory systems of the brain is critical for identifying the neural mechanisms that support SL.

### 1.3 The role of the hippocampus in SL

A central question in the SL literature is whether statistical regularities are extracted by multiple modality-specific systems operating independently or by shared, domain-general regions that act across modalities (Frost et al., 2015), resulting in two competing perspectives on the neural mechanisms underlying SL. One view argues that SL occurs largely outside the hippocampus, relying on modality-specific cortical regions that automatically encode regularities within their respective domains. By this account, hippocampal activity during SL reflects the formation of parallel representations relevant for subsequent memory retrieval but does not drive learning (Batterink et al., 2019; see Conway, 2020, for a review). The second view similarly acknowledges the contribution of modality-specific cortical processing but argues that these sensory representations are subsequently acted upon by shared, domain-general systems, including the hippocampus, which employ a similar computational mechanism across various types of input to support the extraction of statistical regularities and are therefore critical for SL (see Zhou & Turk-Browne, 2025, for a review of hippocampal involvement in SL). Therefore, whether hippocampal involvement reflects an essential component of SL, and how this relates to whether SL is expressed implicitly or explicitly in behaviour, remains an active area of debate in the literature.

#### 1.3.1 Computational and neuropsychological perspectives on hippocampal involvement in SL

One influential account proposing a central role for the hippocampus in SL was put forth by Schapiro et al. (2017). They proposed that the hippocampus accomplishes two competing processes via separate pathways. The trisynaptic pathway, which projects from the entorhinal cortex to the dentate gyrus (DG), CA3, and then CA1 subfields, supports pattern separation by encoding highly similar experiences into distinct, non-overlapping representations. In contrast, the monosynaptic pathway, which projects directly from the entorhinal cortex to CA1, is proposed to support the integration and extraction of regularities from these overlapping experiences, thus supporting SL (**Figure 1**). Using a neural network model that simulated known physiological properties of the hippocampus, they observed that representations of a structured sequence, which could only be learned by tracking transitional probabilities, emerged most strongly in CA1 and could still be accomplished when the trisynaptic pathway was “lesioned”. In contrast, when introducing an explicit cue to the embedded pair boundaries, individual pairs were represented most strongly in the DG and CA3 subfields.

**Figure 1:**
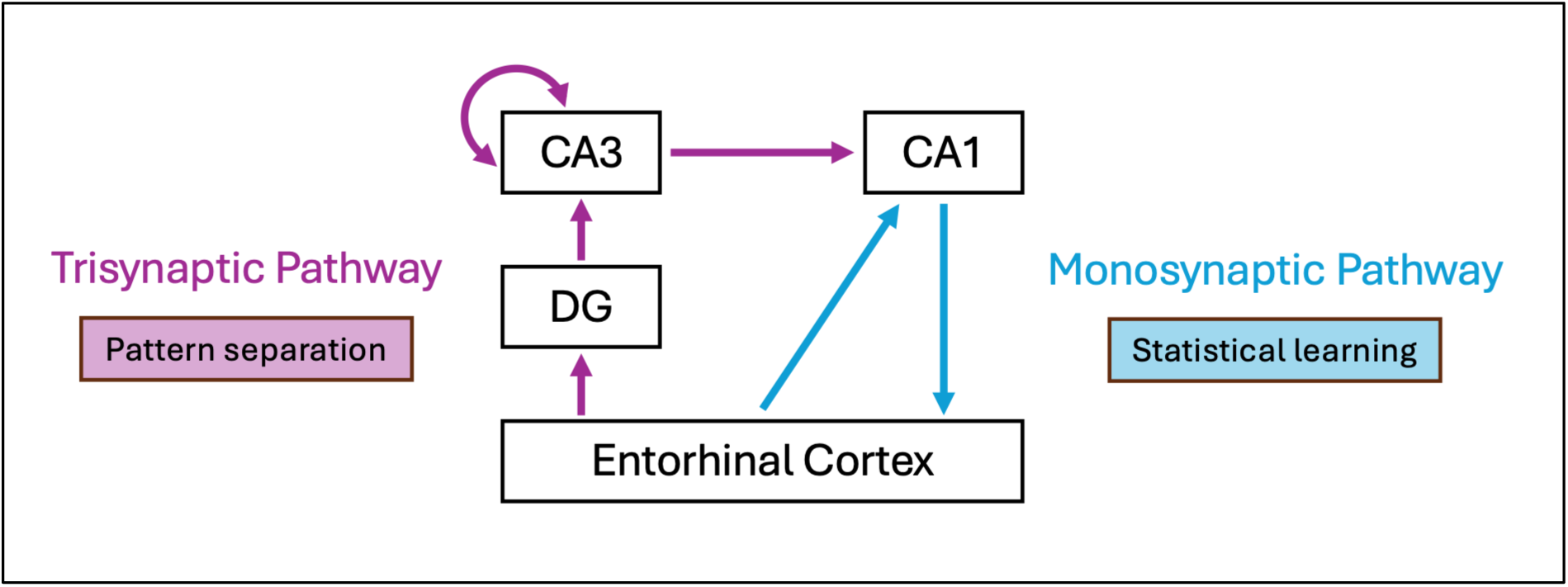
The two pathways of the hippocampus. Adapted from “The prevalence and importance of statistical learning in human cognition and behavior” by B. E. Sherman, K. N. Graves, & N. B. Turk-Browne, *Current Opinion in Behavioral Sciences, 32*, 15-20. https://doi.org/10.1016/j.cobeha.2020.01.015. Copyright 2020 by the authors. CA1/3 = cornu ammonis 1/3, DG = dentate gyrus.

While the trisynaptic pathway, particularly the DG and CA3 subfields, has been consistently implicated in pattern separation in humans (e.g., Bakker et al., 2008; Berron et al., 2016; Lacy et al., 2011; Yassa et al., 2010) and in nonhuman species (e.g., Leutgeb et al., 2007), direct evidence that the monosynaptic pathway contributes to SL remains limited. Empirical studies are imperative to confirm the model’s predictions. Patient-based lesion research, in particular, would provide a unique opportunity to address causal necessity by determining whether disruption of the monosynaptic pathway produces impairments in SL expression.

In contrast to Schapiro et al.’s (2017) computational model, traditional neuropsychological theories of long-term memory have posited that the hippocampus is primarily involved only in explicit or declarative memory systems. According to this framework, explicit memory – including recollection of personal experiences and semantic knowledge – is critically dependent on the medial temporal lobe, including the hippocampus (Squire, 1992). In contrast, implicit or nondeclarative memory processes – including procedural learning, conditioning, nonassociative learning, and priming – rely on a distributed set of structures outside the hippocampus (**Figure 2**; Squire & Zola, 1996). This distinction was largely derived from evidence demonstrating that hippocampal damage can severely impair explicit memory while sparing some forms of implicit memory, whereas damage to other neural systems can produce the opposite pattern. For example, amnesic patients with medial temporal lobe damage performed comparably to healthy controls on an implicit probabilistic classification task despite exhibiting impaired explicit recall for details about the training episode itself (Knowlton et al., 1994). In contrast, patients with Huntington disease and Parkinson disease, which primarily affect corticostriatal systems, were severely impaired on the same implicit probabilistic classification task (Knowlton, Mangels, et al., 1996; Knowlton, Squire, et al., 1996).

**Figure 2:**
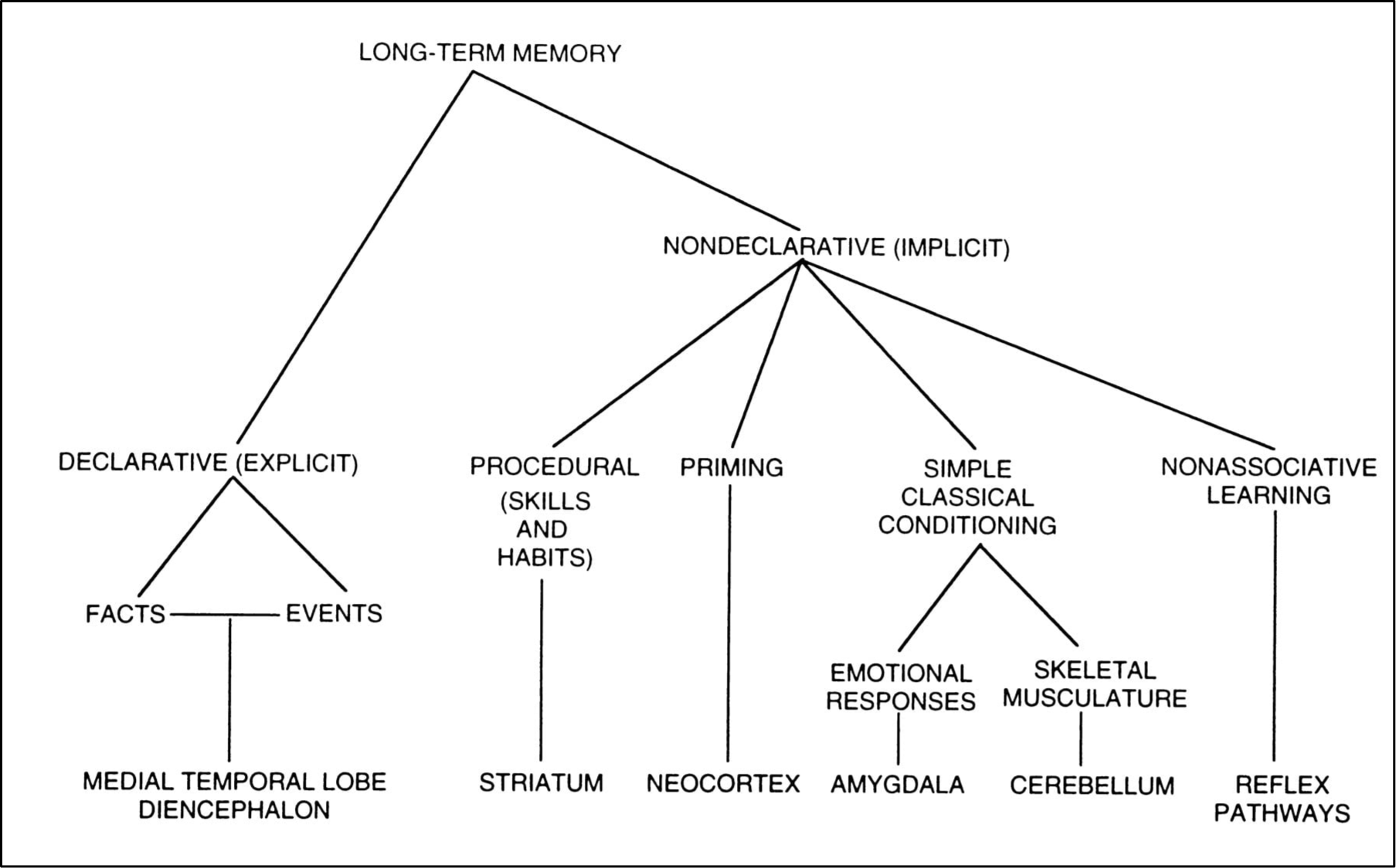
A taxonomy of long-term memory systems together with specific brain structures involved in each system. Reprinted from “Structure and function of declarative and nondeclarative memory systems” by L. R. Squire, & S. M. Zola, *PNAS, 93*(24), 12515-12522. https://doi.org/10.1073/pnas.93.24.13515. Copyright 1996 by the authors.

While this theory has been influential in the memory literature, it has also been challenged by alternative perspectives suggesting that hippocampal contributions are not restricted to explicit or declarative memory, but support relational binding across stimuli to form flexible representations that can be used across memory tasks (Bird & Burgess, 2008; Henke, 2010). These competing perspectives raise important questions regarding whether hippocampal involvement depends on the manner in which learned information is expressed. This distinction has not yet been investigated in the context of SL, largely because many studies on SL have only used explicit memory measures to assess learning after exposure, while the use of implicit measures is relatively more recent. Under this framework, it is possible that the hippocampus is necessary only for the intentional retrieval of stored memory representations required for explicit judgements, but it may not be necessary for the incidental learning or implicit expression of SL. This would add an additional dimension to Schapiro et al.’s (2017) computational model, which currently makes no distinctions about the learning or memory conditions, and therefore implies that the hippocampus is necessary for all aspects of SL.

Although implicit and explicit memory are theoretically distinct constructs, a critical consideration is the extent to which behavioural measures accurately map onto these underlying memory processes. As Jacoby (1991) argued, there is no such thing as a “process-pure” task; rather, performance on any memory measure may reflect varying contributions of automatic and intentional retrieval processes. This challenge highlights the importance of how implicit and explicit memory processes are operationalized. Building on this issue, Schacter et al. (1989) revisited earlier conceptualizations of implicit and explicit memory that emphasized the role of conscious awareness (Schacter, 1987), arguing that retrieval intentionality provides a more useful operational criterion for distinguishing between these memory processes. Similar distinctions have also been considered within neuropsychological models of hippocampal contributions to memory. For example, Leritz et al. (2006) proposed that behavioural measures may fall along a continuum ranging from highly explicit, hippocampally dependent processes, such as spontaneous recall, to highly implicit, hippocampally independent processes, such as repetition priming, with many tasks relying on varying contributions from both systems.

This framework has important implications for behavioural assessments of SL. Although both the familiarity rating and forced-choice recognition tasks require explicit responses, they may differ in the extent to which they rely on intentional retrieval of learned information. Familiarity ratings of the kind employed in SL research require participants to evaluate each item independently and may therefore rely on retrieval of highly precise memory representations of the learned sequences (Wang, Rosenbaum, et al., 2023). In contrast, forced-choice recognition places fewer demands on the retrieval of such precise representations, as judgements can be based on the relative familiarity between competing alternatives (Smalle & Boagaerts, 2024; Voss & Paller, 2008; Wang, Rosenbaum, et al., 2023). Consistent with limitations in explicit access to knowledge acquired through incidental SL, participants are often unable to explicitly verbalize the statistical regularities they have acquired despite demonstrating learning on recognition-based measures (Endress, 2024; Perruchet & Pacton, 2006). In contrast, the target detection task places no overt demands on explicit retrieval. Since responses are made online during rapid stimulus presentation, participants have little opportunity to deliberately retrieve previously learned sequences, making performance more likely to reflect automatic priming and/or prediction. Furthermore, since explicit testing may influence subsequent expression of implicit knowledge, the target detection task is typically administered before explicit memory measures so that isolated presentations of words do not provide additional word-boundary cues (Batterink et al., 2015; Turk-Browne et al., 2005).

Accordingly, these three behavioural tasks provide promising proxies for examining implicit and explicit dissociations in the context of SL expression, although they may not provide pure indices of implicit or explicit memory. Evidence supporting separate implicit and explicit memory systems has largely come from single dissociations observed across different experimental tasks, often involving different materials and cognitive demands (Leritz et al., 2006; Squire & Zola, 1996). In contrast, SL paradigms have the advantage that implicit and explicit expressions of the same learned regularities can be assessed within a single experimental framework. Differences across behavioural measures can therefore be more directly attributed to the memory processes supporting expression, rather than differences between the types of stimuli or the conditions under which they were learned.

#### 1.3.2 Evidence linking the hippocampus to SL

Investigations of hippocampal involvement in SL have produced mixed findings, likely reflecting substantial methodological differences across studies. Previous work has employed a wide range of approaches, including functional Magnetic Resonance Imaging (fMRI; Ellis et al., 2021; Schapiro et al., 2012; Sherman & Turk-Browne, 2020; Turk-Browne et al., 2009), intracranial recordings (Henin et al., 2021; Herrera-Chaves et al., 2026; Ramos-Escobar et al., 2022; Tacikowski et al., 2024), and lesion studies (Covington et al., 2018; Schapiro et al., 2014; Wang, Rosenbaum, et al., 2023), using different stimulus modalities, presentation rates, and markers of learning.

Overall, slower-paced visual paradigms using fMRI have generally implicated the hippocampus in the extraction and integration of statistical regularities, reporting increased hippocampal activation (Ellis et al., 2021; Turk-Browne et al., 2009), representational changes (Schapiro et al., 2012), and predictive responses (Sherman & Turk-Browne, 2020). In contrast, rapid auditory paradigms using intracranial EEG have more commonly observed cortical neural entrainment to the temporal structure of the input, with less consistent evidence for hippocampal involvement.

These intracranial studies report mixed findings, with some observing a peak in hippocampal power at the structured frequency (Ramos-Escobar et al., 2022) while others do not observe reliable entrainment in the hippocampus (Henin et al., 2021; Herrera-Chaves et al., 2026). As argued by Zhou and Turk-Browne (2025), the variability observed across studies is likely driven by differences in modalities, stimulus materials, and stages of learning that are captured by these methodological approaches.

While these approaches provide important insights into the neural mechanisms engaged during the learning process itself, they cannot provide evidence concerning whether the hippocampus plays a causal role in SL and its different expressions in behaviour. Much of the work on this relationship uses a lesion-based approach.

Schapiro et al. (2014) provided one of the first demonstrations that hippocampal damage can impair behavioural performance in SL in a single case study of an amnesic patient with a bilateral hippocampal lesion and additional medial and anterior temporal lobe damage. They exposed this participant to four versions of a SL paradigm using different stimulus types across auditory and visual modalities, including scenes, shapes, syllables, and tones, then assessed learning using a forced-choice recognition task. The patient performed at or below chance across all versions and performed significantly worse than healthy controls. In a subsequent study, Covington et al. (2018) extended this work by examining four amnesic patients with varying degrees of hippocampal damage and one patient with more extensive medial temporal lobe damage. Across the same four versions of the SL paradigm, patients performed significantly worse than healthy controls on the forced-choice recognition tasks, although some individuals demonstrated above-chance learning, suggesting that SL remained possible but was reduced following hippocampal damage. Since both studies only used a single explicit measure to assess learning and did not include any implicit measures of learning, this leaves open the possibility that the patients’ knowledge of the streams’ structures could have been expressed implicitly despite showing impaired performance on the explicit recognition measure.

To date, only one study has directly tested Schapiro et al.’s (2017) computational model using both explicit and implicit measures of SL. Wang, Rosenbaum, et al. (2023) conducted a case study on patient BL, an amnesic patient with a highly selective bilateral lesion to the dentate gyrus that spared the rest of the hippocampus (Baker et al., 2016; Kwan et al., 2015), primarily affecting the trisynaptic pathway, but likely not the monosynaptic pathway. BL demonstrated impaired performance on the explicit familiarity rating task, while his performance on the implicit target detection task remained comparable to age-matched healthy controls. BL also demonstrated impaired performance on auditory (Wang, Rosenbaum, et al., 2023) and visual (Baker et al., 2016) versions of the Mnemonic Similarity Task (Stark et al., 2015), used to probe pattern separation. This confirmed the model’s predictions about the trisynaptic pathway, and more specifically the dentate gyrus, supporting pattern separation in a domain-general fashion.

While the findings from these studies generally support that hippocampal damage can disrupt the explicit expression of SL, there is still little evidence to address whether the hippocampus contributes to the implicit expression of SL. The first two studies only assessed learning using a single explicit measure, failing to capture all aspects of knowledge expression. They also involved patients with extensive hippocampal or medial temporal lobe damage, making it impossible to determine the contribution of individual hippocampal subfields, even in the case of explicit SL expression (Covington et al., 2018; Schapiro et al., 2014). Conversely, the selective dentate gyrus lesion observed in patient BL provides evidence regarding the role of one hippocampal subfield (Wang, Rosenbaum, et al., 2023) but cannot determine whether the intact implicit expression of SL observed in this case reflects the engagement of other preserved subfields and neural communication through the monosynaptic pathway, or the contributions of extra-hippocampal brain regions. Therefore, further research is needed to determine how variable hippocampal pathology affects different expressions of SL.

We extend these lesion-based studies of SL to patients with temporal lobe epilepsy (TLE). This population provides a unique model for investigating the contribution of hippocampal integrity to SL since TLE is often characterized by variable patterns of hippocampal pathology accompanied by broader memory impairments.

### 1.4 Temporal lobe epilepsy

Epilepsy is a neurological disorder characterized by a predisposition to recurrent, unprovoked seizures resulting from abnormal, excessive, or synchronous neuronal activity in the brain (Fisher et al., 2014). Seizures can vary widely in their clinical presentation depending on the brain regions involved, but may include changes in awareness, sensory experiences, emotional states, autonomic responses, and involuntary movements (Vinti et al., 2021). Epilepsy is relatively common, affecting approximately 52 million people worldwide as of 2021 (GBD Epilepsy Collaborators, 2025). It can arise from a diverse range of causes, including genetic predispositions, developmental abnormalities, structural brain lesions, infections affecting the central nervous system, and metabolic disturbances (Falco-Walter et al., 2018). Although many patients with epilepsy achieve seizure control with antiseizure medications, approximately one-third continue to experience seizures, a condition referred to as “drug-resistant” epilepsy (Kwan et al., 2010). Therefore, patients with this type of epilepsy often undergo surgical evaluation, as resection or ablation of the epileptogenic tissue may provide effective seizure control (Kanner et al., 2022; Nouby et al., 2026).

TLE is the most common form of focal epilepsy, in which seizures originate from within the temporal lobe, most frequently involving medial temporal lobe structures such as the hippocampus (Bertram, 2009; Engel, 2001). In TLE, the seizure onset zone is typically unilateral, with structural pathology most pronounced ipsilateral to (i.e., on the same side as) the seizure onset zone, although bilateral cases do occur (Diehl & Lüders, 2000).

#### 1.4.1 Hippocampal abnormalities in TLE

The most common pathology observed in drug-resistant TLE is hippocampal sclerosis, which is characterized by selective neuronal loss and gliosis within the hippocampus (Blümcke et al., 2013; Curia et al., 2014). Rather than producing uniform damage, TLE preferentially affects specific hippocampal subfields. Histopathological and high-resolution structural MRI studies have consistently demonstrated significant volume reduction in the hippocampus that is ipsilateral to the seizure onset zone, with the greatest neuronal loss observed in the CA1 subfield in some studies, with some implications of CA2, CA3, CA4, dentate gyrus, and subiculum (e.g., Blümcke et al., 2013; Kim et al., 2015).

Unfortunately, hippocampal abnormalities in TLE are not always detectable using conventional clinical MRI methods, resulting in a subset of cases classified as “MRI-negative” (Gill et al., 2025). Yet, these patients often exhibit subtle hippocampal alterations when examined using quantitative volumetric and subfield-specific analyses (Ellsay & Winston, 2024; Ripart et al., 2024). Furthermore, histopathological examination following surgical resection has shown that approximately 10 percent of MRI-negative focal epilepsy cases demonstrate hippocampal sclerosis despite the absence of visible abnormalities on clinical imaging (Wang et al., 2013). Together, these findings suggest that hippocampal pathology in TLE is not restricted to cases with MRI-visible lesions and may vary in severity across patients.

#### 1.4.2 General memory abilities in TLE

Memory abilities in patients with TLE are typically assessed as part of a comprehensive neuropsychological evaluation conducted within clinical contexts. These assessments are used to characterize memory deficits as a clinical symptom of their neurological disorder, inform surgical planning, and monitor cognitive outcomes following treatment (Baxendale, 2020). Standardized neuropsychological tests predominantly assess explicit memory by requiring participants to encode and intentionally retrieve information through various tasks involving free or cued recall and recognition. Performance is typically assessed both immediately after learning to assess short-term memory and again following a delay of approximately 20 to 30 minutes to assess longer-term retention (e.g., Lezak et al., 2012; Wechsler, 2009). Consequently, studies investigating memory in clinical populations have largely relied on explicit neuropsychological measures, whereas implicit memory has been examined far less frequently and primarily within experimental research settings.

TLE is consistently associated with impairments in explicit memory, particularly for tasks requiring the encoding and conscious retrieval of verbal information. Across neuropsychological studies, patients with TLE perform worse than both healthy controls and patients with other forms of epilepsy, with one of the most robust findings being impaired verbal episodic memory in Left TLE. These deficits are commonly observed on tasks involving story learning and word-list recall and recognition, consistent with the specialization of the left hemisphere in linguistic processing (Lacritz et al., 2004; Neudorf et al., 2020). In contrast, Right TLE is more commonly associated with impairments in nonverbal and visuospatial memory, including tasks involving facial recognition and spatial navigation, reflecting the contribution of the right hemisphere to perceptual and spatial processing (Abrahams et al., 1997; Meletti et al., 2003). In Bilateral TLE, where epileptic activity affects both temporal lobes, memory impairments are typically more severe and widespread, reflecting reduced capacity for compensation by the contralateral hemisphere (Baggio et al., 2023).

In contrast, evidence for implicit memory functioning in TLE is more variable. Some studies have reported impairments on implicit tasks, particularly when they involve verbal material or place greater demands on relational processing and conceptual knowledge (Savage et al., 2002; Zaidel et al., 1994; Zaidel et al., 1998). However, interpretation of these findings is complicated by methodological factors, including dissociations derived from different stimulus materials and tasks, as well as the order of task administration, with implicit performance potentially contaminated when explicit tasks are administered first (Blaxton, 1992; Savage et al., 2002; Zaidel et al., 1994; Zaidel et al., 1998). In contrast, studies that more directly examined dissociations between implicit and explicit memory by assessing the same material under different retrieval conditions have generally demonstrated preserved implicit memory despite impairments in explicit memory (Bilingsley et al., 2002; Del Vecchio et al., 2004). For example, patients with TLE have shown intact priming behavioural effects on word identification and word generation tasks, even when explicit recall and recognition for the same material were significantly impaired (Billingsley et al., 2002). Overall, this literature suggests a robust and relatively selective impairment of explicit memory in TLE, particularly for verbal material and left temporal pathology, while more research is needed to investigate the effects of TLE on implicit memory processes.

#### 1.4.3 SL expression in TLE

To date, only one other study that we are aware of has investigated the impact of TLE on behavioural performance in SL tasks. Aljishi et al. (2024) used a visual SL paradigm in which participants viewed a continuous stream of nonlinguistic hieroglyph symbols organized into deterministic pairs. Following exposure, participants’ knowledge of the statistical structure was assessed using an association test in which they were presented with the first symbol of a pair and asked to select the second corresponding symbol from two alternatives. Overall, patients with epilepsy (*n* = 38) performed above chance and did not differ significantly from healthy controls (*n* = 28). Seizure onset zone – temporal (*n* = 23) versus extratemporal (*n* = 15) – was not associated with SL performance, although the extratemporal lobe epilepsy group performed below chance. With respect to hippocampal integrity, CA1 and CA2/3 subfield volumes reliably predicted SL performance, with CA1 volume negatively correlating, and CA2/3 volume positively correlating, with performance (Aljishi et al., 2024).

Taken together, these findings provide mixed evidence regarding the role of the hippocampus in SL, as the structural findings are difficult to interpret. Specifically, reduced CA2/3 volume was associated with poorer SL performance, consistent with the expectation that reduced hippocampal integrity leads to impaired SL. In contrast, reduced CA1 volume was associated with better SL performance, which is difficult to reconcile within this account. Similarly, the absence of an association between seizure onset zone and SL performance is unexpected given that epileptic activity in the temporal lobe typically affects the ipsilateral hippocampus and hippocampal lesions have led to impairments in SL expression (Blümcke et al., 2013; Covington et al., 2018; Diehl & Lüders, 2000; Kim et al., 2015; Schapiro et al., 2014; Wang, Rosenbaum, et al., 2023).

While these findings may provide preliminary evidence for hippocampal involvement in SL, several important questions remain. First, because the paradigm was entirely visual, it is unclear whether these findings generalize to auditory SL, as previous research has shown that SL is subject to modality-specific constraints (Conway & Christiansen, 2005) and that individual differences in SL show little correspondence across these two modalities (Siegelman & Frost, 2015). Although the hippocampus has been proposed to function as a domain-general hub for SL (Frost et al., 2015), this hypothesis has largely been supported by studies employing only explicit measures of SL (Covington et al., 2018; Schapiro et al., 2014). In their study, Aljishi et al. (2024) assessed learning exclusively using an explicit association test, which is conceptually similar to the forced-choice recognition tasks used previously to measure explicit expression of SL. Consequently, it remains unclear whether hippocampal involvement extends to the implicit expression of SL.

Together, these findings establish TLE as a valuable model for investigating hippocampal contributions to SL expression, while highlighting an important gap in the literature: whether hippocampal involvement is limited to the explicit expression of SL or extends to both explicit and implicit expressions.

### 1.5 The current study

The current study aimed to investigate whether TLE differentially affects implicit and explicit expressions of SL. We presented 27 patients with TLE and 34 healthy controls with a continuous auditory speech stream that contained four embedded trisyllabic words, based on the paradigm used by Wang, Rosenbaum, et al. (2023). Following exposure, learning was assessed implicitly using a target detection task and explicitly using a familiarity rating task and a two-alternative forced choice (2AFC) recognition task.

Given previous work on the heterogeneous nature of hippocampal pathology in TLE (e.g., Blümcke et al., 2013; Kim et al., 2015) and previous findings that individuals with hippocampal lesions demonstrate impaired performance on explicit measures of SL compared to healthy controls (Covington et al., 2018; Schapiro et al., 2014; Wang, Rosenbaum, et al., 2023), we predicted that patients with TLE would similarly show reduced performance on our explicit measures of SL. Regarding implicit expression of SL, there are two alternative hypotheses. If the hippocampus contributes to SL in general, as proposed by Schapiro et al.’s (2017) computational model, then TLE patients should also exhibit impaired performance on our implicit measure of SL. Alternatively, if the implicit expression of SL relies primarily on structures outside the hippocampus, consistent with traditional neuropsychological models of long-term memory (Squire & Zola, 1996), then performance on our implicit measure of SL should be relatively preserved despite hippocampal pathology.

## 2 Methods

### 2.1 Participants

A total of 41 patients with drug-resistant, MRI-negative epilepsy and 36 neurotypical, healthy controls were recruited to participate in the study. All participants provided informed consent prior to participation in accordance with the Research Ethics Board at the University of Western Ontario.

Patients were recruited through *University Hospital* in London, ON, Canada, where they were admitted to the *Epilepsy Monitoring Unit* to undergo stereoencephalography for seizure onset localization. Patients were excluded from the final analysis if they did not complete the entire study (*n* = 2) or if their epilepsy diagnosis did not include a clinical TLE classification determined by hospital clinicians using established diagnostic criteria and clinical judgement (*n* = 12). Demographic information of all patients in the final sample is included in **Table 1**.

**Table 1:** Patient demographics.

| Patient | Age | Sex | Education | Handedness | TLE<br>Classification |
| --- | --- | --- | --- | --- | --- |
| 1 | 25 | F | 12 | Right | Right |
| 2 | 19 | F | 11 | Right | Right |
| 3 | 39 | M | 16 | Left | Left |
| 4 | 34 | F | NR | Right | Right |
| 5 | 35 | F | NR | Right | Left |
| 6 | 24 | F | 12 | Right | Right |
| 7 | 36 | M | 15 | Right | Left |
| 8 | 27 | F | NR | Right | Unspecified |
| 9 | 25 | M | NR | Right | Left |
| 10 | 26 | M | 13 | Right | Bilateral |
| 11 | 25 | M | 12 | Left | Left |
| 12 | 39 | M | 12 | Right | Left |
| 13 | 51 | M | 12 | Right | Bilateral |
| 14 | 36 | M | 10 | Right | Bilateral |
| 15 | 44 | F | 12 | Right | Bilateral |
| 16 | 46 | F | 13 | Left | Bilateral |
| 17 | 54 | M | 12 | Right | Bilateral |
| 18 | 26 | M | 14 | Right | Bilateral |
| 19 | 23 | F | 12 | Left | Left |
| 20 | 37 | M | 18 | Right | Left |
| 21 | 29 | F | 14 | Right | Bilateral |
| 22 | 47 | F | 16 | Left | Right |
| 23 | 31 | F | 12 | Right | Right |
| 24 | 30 | M | 16 | Right | Left |
| 25 | 22 | F | 14 | Right | Left |
| 26 | 32 | F | 18 | Right | Right |
| 27 | 58 | F | 16 | Right | Left |

*Note.* Age and education are reported in number of years. F = female, M = male, NR = not reported, TLE = temporal lobe epilepsy.

Healthy controls were recruited through Western’s BrainsCAN registry, community advertisements, and an additional research protocol in which participants had consented to be contacted again for future studies. All participants were required to be fluent in English and to have normal or corrected-to-normal hearing and vision. Native English-speaking participants were additionally required to pass the Montreal Cognitive Assessment (MoCA), a validated screening tool for mild cognitive impairment (Nasreddine et al., 2005). However, performance on the MoCA has been shown to be influenced by linguistic and cultural factors, raising concerns regarding the validity of scores across diverse populations (O’Driscoll & Shaikh, 2017). Thus, to avoid false-positive identification of cognitive impairment in individuals for whom English is a second language, the MoCA version 8.1 English was used as an exclusion criterion only for participants who identified English as their first language. Two participants fell below the cut-off for normal range of performance (i.e., a score of less than 26 out of 30) and were therefore excluded from the final analysis.

Following exclusions, the final sample consisted of 27 TLE patients between the ages of 19 and 58 years (15 females; average age = 34.1 years; average years of education = 13.6; 22 right-handed) and 34 healthy controls between the ages of 19 and 52 years (16 females; average age = 30.9 years; average years of education = 17.3; 32 right-handed). As summarized in **Table 2**, TLE patients did not significantly differ from controls in terms of age (*t*(59) = -1.18, *p* = .244), sex (χ^2^ = .44, *p* = .51), nor handedness (χ^2^ = 2.37, *p* = .124). They did, however, significantly differ in education (*t*(55) = 4.99, *p* < .001), with the TLE patient group being less educated on average.

**Table 2:**
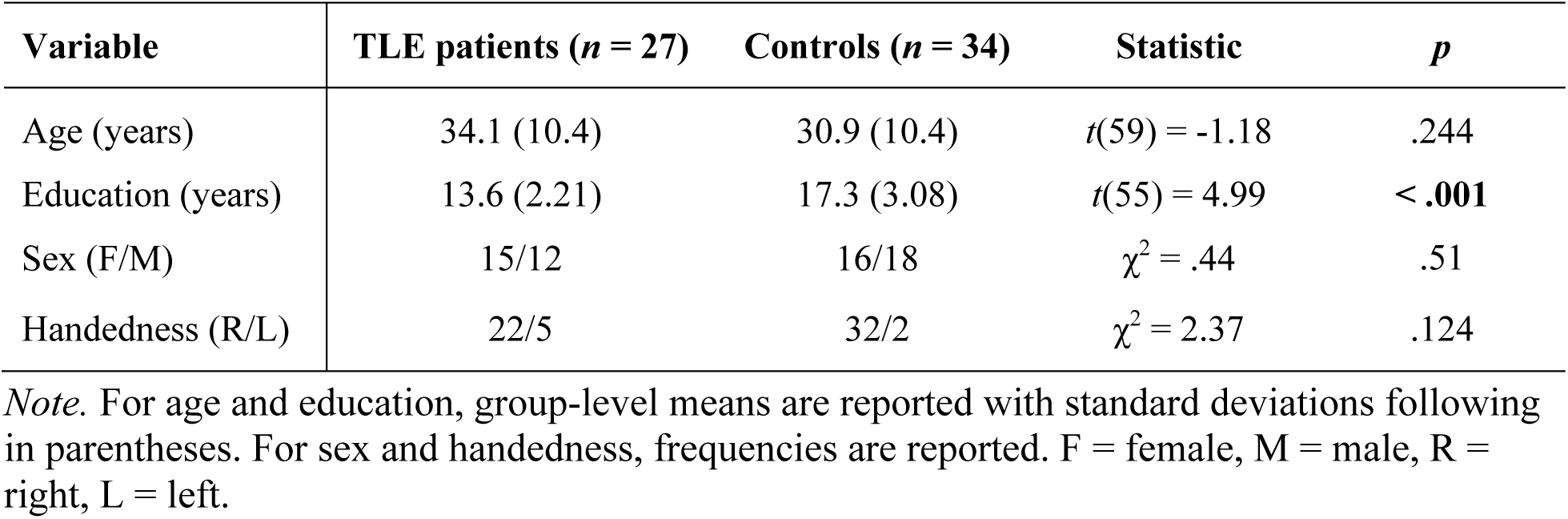
Sample characteristics.

### 2.2 Stimuli

#### 2.2.1 Statistical learning paradigm

Stimuli for this auditory SL paradigm were adapted from Batterink & Paller (2019) and consisted of 12 distinct syllables that followed a consonant-vowel format (e.g., *ba*). Syllables were recorded by a male native English speaker with neutral intonation and no co-articulation between syllables, then spliced into individual audio files ranging from 220 to 250 ms in duration. The syllables were combined to form four trisyllabic nonsense words (*bafuko, fetisu, regeme, rupuni*) with each syllable consistently occurring in either the first, second, or third position. For simplicity, these stimuli will be referred to as “words”.

##### 2.2.1.1 Structured exposure phase

A continuous audio speech stream was created by concatenating the four words in a pseudorandom order at a fixed rate of 300 ms per syllable. The order remained consistent across participants. There were no explicit word boundaries (e.g., pauses, cues) to indicate syllable groupings. Each word was repeated 90 times with the constraint that the same word could not follow itself consecutively, resulting in a total stream duration of 5 mins 24 s.

##### 2.2.1.2 Target detection task

A total of 36 shorter speech streams were created in a similar fashion to the Exposure Phase speech stream, except the four words were only repeated four times each for a total duration of 14.4 s. The 36 streams were divided into three blocks of 12, with each stream containing one designated target syllable. Each of the 12 syllables served as the target once per block, with first, second, and third position targets equally distributed across the 12 streams. The order of targets was randomized between blocks to minimize potential trial order effects. Within each stream, the target syllable appeared four times with the constraint that the target’s designated word could not occur as either the first or last word of the stream. Across the task, each syllable served as a target 12 times, yielding 48 target presentations in each triplet position, and a grand total of 144 target presentations overall.

Two practice streams were designed in a similar way but using nine different syllables than those included in the rest of the paradigm and were also recorded using a different speech synthesizer voice. The syllables were combined to form three trisyllabic nonsense words (*padoti, tupelo, widaku)* that were each repeated five times for a total duration of 13.5 s. Thus, each practice stream contained five targets.

##### 2.2.1.3 Familiarity rating and two-alternative forced-choice recognition tasks

The 12 syllables were recombined to form three different word types: (1) Words, consisting of the original configurations from the exposure phase, (2) Partwords, in which two consecutive syllables from an original Word were combined with the remaining position syllable from another Word (e.g., *ba + puni, rege + su*), and (3) Nonwords, in which all three syllables were recombined from different Words (e.g., *ni + su + ba*).

#### 2.2.2 Perceptual similarity task

Stimuli for this task were adapted from Wang, Rosenbaum, et al. (2023) and consisted of 25 distinct sounds that are common in the environment, including sounds produced by humans, animals, and man-made objects (instruments, household, and outdoor). Five of these sounds were repeated to form five ‘Identical’ sound pairs while another five were paired with a similar ‘Lure’ sound from the same semantic category (e.g., two different “cat meowing” sounds) to create five ‘Similar’ sound pairs. The remaining 10 sounds were grouped into five ‘Different’ sound pairs, with each pair selected from within the same semantic category (e.g., two instrument sounds: a trumpet and a cello). Altogether, this resulted in 15 trials, with five trials per pair type (Identical, Similar, Different) and each category (animal, human, instrument, household, and outdoor) represented once within each pair type.

### 2.3 Experimental procedure

Informed consent was obtained from all participants prior to testing. Healthy controls provided consent directly to the experimenter, whereas patient consent was obtained by a trained research assistant from the patient’s clinical care team, in accordance with the hospital’s research procedures and the ethics protocol guiding the study.

Following consent, patients completed tasks on a desktop computer that was wheeled to their bedside. A single monitor was positioned in front of them at a comfortable reading distance for instructions. They were given a pair of JBL TUNE 110 earbuds that they were permitted to keep after the experiment. Earbuds were used to minimize external noise and distractions, as patients were assigned to a bed within a common area that could include up to 11 other patients, clinical staff, and visitors present at any given time. Responses were collected via a wireless keyboard where necessary. As compensation, patients received an Amazon gift card at a rate of $20 CAD per hour, delivered via email when possible. If they were unable to provide a valid email address, they were handed a physical gift card.

Healthy control participants completed the tasks on a MacBook Air laptop with a 15-inch screen display, placed at a comfortable reading distance for instructions. They were provided with over-ear headphones to use for the duration of the experiment. These headphones were sanitized after each use and reused across participants. Headphones were used to minimize disruptions from other testing sessions being conducted in adjacent rooms. Participants made key responses where necessary using the laptop’s built-in keyboard. As compensation, they were paid at a rate of $14 CAD per hour.

Before beginning the experiment, the experimenter verbally gave a general overview of the experimental procedure, encouraged them to ask questions for clarification, and reminded them that they could terminate their participation at any point during the study. Participants first completed a hearing check to ensure that the audio could be heard clearly and at a comfortable volume level. The experimenter adjusted the volume, as needed. Prior to each task, participants were given detailed instructions for the task on the computer monitor for them to read.

Task administration varied across participants due to modifications made to the procedure over the course of data collection. The first cohort of patients (*n* = 16) was initially recruited for another study to primarily obtain intracranial recordings during SL task performance. These patients completed only the structured exposure phase, target detection task, familiarity rating task, and random exposure phase. Subsequent patients (*n* = 11) recruited for this study additionally completed the SL two-alternative forced-choice task and the perceptual similarity task, while the SL random exposure task was removed. A pattern separation paradigm, including mnemonic similarity tasks for auditory stimuli was also administered as part of another independent research project. For this second group, a 30-minute break was included between the SL paradigm and subsequent tasks to mitigate fatigue. Healthy controls were given the latest version of the task battery, with the exception that they also completed a demographic questionnaire and the MoCA during the 30-minute break period. A schematic overview of the different task versions administered across participants is shown in **Figure 3**. Overall, behavioural testing lasted approximately one to two hours, depending on the version administered. Two participants in the healthy control group withdrew from the study after completing the MoCA.

**Figure 3:**
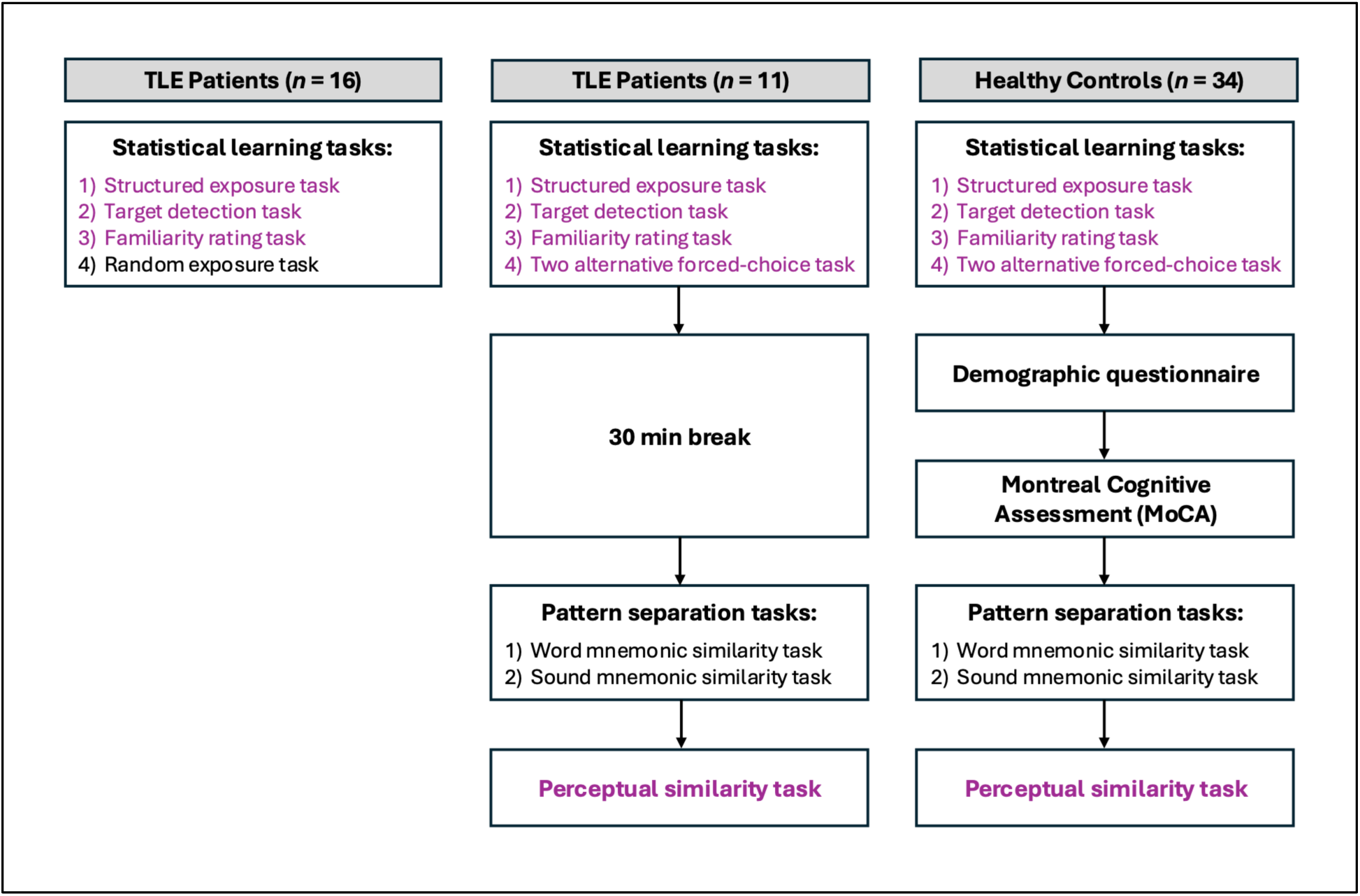
Variations in experimental procedure across participants. The first group of TLE patients (*n* = 16) were tested only on a set of statistical learning tasks, which included a random exposure task. Later, the procedure changed for all subsequent TLE patients (*n* = 11) such that the random exposure task was replaced with the two-alternative forced-choice task, and the addition of two pattern separation tasks and a perceptual similarity task after a 30-minute break. Healthy controls (*n* = 34) received a similar procedure as the second group of patients except they completed a demographic questionnaire and the Montreal Cognitive Assessment during the 30-minute break period. The data from the tasks highlighted in purple are being analyzed as part of this study.

All tasks were either identical to or adapted from the tasks used in Wang, Rosenbaum, et al. (2023). They were programmed and presented using *PsychoPy*, with the hospital’s desktop running version 2022.2.4 and the lab-based laptop running version 2022.2.5 (Peirce et al., 2019).

#### 2.3.1 Statistical learning paradigm

##### Structured exposure phase

This task served as a learning phase for the statistical regularities that would later be probed using implicit and explicit memory measures. Participants were instructed to listen to an artificial speech stream under the premise that it was an “alien” language that the researchers needed help deciphering. They were not told about the embedded structure of the speech, just to listen passively for the entire duration without any breaks or cover tasks. A schematic of the structured exposure phase is shown in **Figure 4A**.

**Figure 4:**
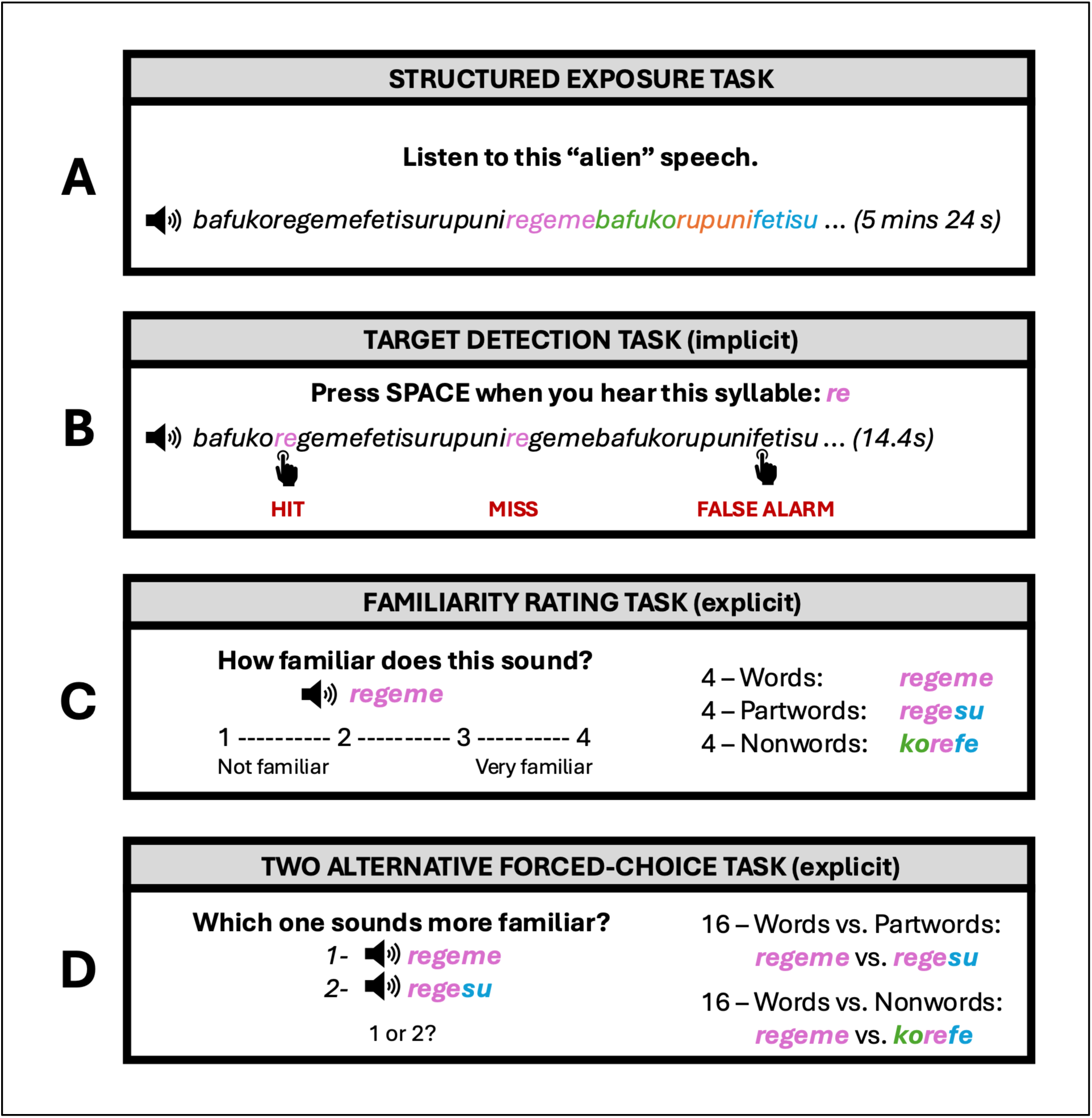
Statistical learning task design and procedure. (A) Structured exposure task – All participants first listened to a continuous speech stream with an embedded structure of repeating trisyllabic words for 5 minutes and 24 seconds. (B) Target detection task – Participants then listened to shorter versions of the structured exposure stream and were instructed to make a key response when they heard a specific syllable. Hits are responses made within 1200ms of the target onset. Misses are target syllables without a response. False alarms are responses made outside the 1200ms target window. (C) Familiarity rating task – Participants were asked to rate the familiarity of three item types (word, partword, nonword) on a scale from 1 (not familiar) to 4 (very familiar). (D) Two-alternative forced-choice task – A subset of patients (*n* = 11) and all healthy controls were presented with two items (a word and either a partword or nonword) and asked to choose the one that sounded the most familiar.

##### Target detection task

This task served as an implicit measure to probe knowledge acquired through SL. Participants were given a specific target syllable to detect during shorter streams of the “language” presented in the initial exposure phase. At the beginning of each trial, participants saw the target syllable written on the screen (e.g., “ba”) and listened to the individual syllable recording twice. Participants were instructed to press the spacebar as quickly and as accurately as possible whenever they heard the target syllable during the trial audio. The written form of the syllable remained on the screen throughout the trial to prevent forgetting during the task. This task is depicted in **Figure 4B**.

Participants completed two practice trials prior to the 36 test trials that used a different voice and syllable set. The practice trials allowed participants to get used to the task by receiving feedback on their performance. Specifically, the number of target syllables detected (hits) and average reaction time (RT; in seconds) were displayed on the screen after each practice trial. Participants were warned that they would not receive any feedback during the test trials. They were also told that they could complete the task at their own pace and were encouraged to take short breaks in between trials as needed.

This implicit measure aimed to assess participants’ knowledge of the structured exposure phase through behavioural expression that does not require the conscious or intentional retrieval of stored information from memory. If participants learned the embedded statistical regularities, then they should be able to predict later (i.e., second and third) position syllables from the initial (i.e., first and second) position syllables. Behaviourally, this should facilitate faster RTs across positions (first > second > third; Batterink & Paller, 2017; Batterink et al., 2019).

##### Familiarity rating task

This task served as an explicit measure to probe knowledge acquired through SL. The experimenter informed participants that there were embedded “words” in the audio, such that there were syllables that always occurred together and repeated many times throughout. Participants were told that they would hear a “word” that may or may not have been in the original audio and were instructed to rate its familiarity on a scale from 1 (“Not familiar”) to 4 (“Very familiar”). Across the 12 trials, participants listened to each of the four Words, four Partwords, and four Nonwords, presented in a pseudorandom order that was counterbalanced across participants to minimize order effects. Participants were warned prior to beginning the task that each item would only play once so they should listen carefully. A schematic of the familiarity rating task is shown in **Figure 4C**.

This explicit measure required participants to intentionally retrieve stored representations from their memory to make familiarity judgements about the presented item. If they learned the triplet word units, then they should rate Words as being the most familiar, followed by Partwords, and Nonwords being the least familiar (Batterink & Paller, 2017; Batterink et al., 2019)

##### Random exposure phase

This task served as a control condition for a separate research question and is therefore not being analyzed as part of this study. A different set of 12 syllables were repeated to form a continuous speech stream but it had no underlying statistical structure. Similar to the structured exposure, patients listened to it passively for approximately five minutes without breaks or a cover task.

##### Two-alternative forced-choice task

This task served as an additional explicit measure to probe knowledge acquired through SL. Participants were told they would hear two different “words” and were instructed to choose the one that sounded most familiar to them by pressing 1 (first word was more familiar) or 2 (second word was more familiar) on the keyboard. Across the 32 trials, each of the four Words was paired with each of the four Partwords and four Nonwords, with Words appearing first and second in the queue at an equal rate. The 32 trials were divided into four blocks of eight, in which each unique Partword and Nonword would appear once in a random order before they could be repeated. Like the familiarity rating task, participants were warned prior to beginning the task that the two items would only play once with no option to repeat. This task is depicted in **Figure 4D**.

This explicit measure required participants to intentionally retrieve stored representations from their memory to make recognition judgements regarding the two presented items. If they learned the triplet word units, then they should choose Words as being more familiar than both the Partword and Nonword foils.

#### 2.3.2 Demographic questionnaire

After completing the SL paradigm, all participants were given a 30-minute break to rest from the computerized tasks. During this time, healthy controls completed a paper-based demographic questionnaire.

Participants were asked to provide general demographic information, as well details about their language background, neurological history, general sensory abilities, and current cognitive state.

#### 2.3.3 Montreal Cognitive Assessment

During the 30-minute break, healthy controls also completed the paper version of the MoCA. It is a brief, 10-minute screening tool that evaluates eight cognitive domains in the following order: visuospatial/executive functioning, naming, memory, attention, language, abstraction, delayed recall, and orientation (Nasreddine et al., 2005). Scores range from 0 to 30, with scores of 26 or higher considered within the normal range. Consistent with standard administration guidelines, participants with 12 years of education or less received one additional point added to their total score.

#### 2.3.4 Pattern separation paradigm

Data from these tasks are also not being analyzed as part of this study but were included in later revisions of the testing procedure to address a separate research question. Participants completed two different auditory versions of the continuous Mnemonic Similarity Task (MST; Stark et al., 2015): one using artificial linguistic stimuli and another using common environmental sounds (Wang, Rosenbaum, et al., 2023).

#### 2.3.5 Perceptual similarity task

Participants were told they would hear pairs of environmental sounds taken directly from the Sound MST they had just completed. They were instructed to rate how similar the sounds were to each other on a scale from 1 (“Different”) to 4 (“Identical”). Across the 15 trials, five pairs each were Identical, Similar, and Different. The pairs were presented in a pseudorandom order, counterbalanced across participants to minimize order effects. Prior to beginning the task, participants were warned that the two sounds would only play once and were instructed to listen carefully.

This task was included to assess auditory perceptual abilities, as patients with TLE have been shown to exhibit deficits in auditory processing and sound discrimination (Angeli et al., 2024). If auditory perception is intact, then patients should be able to discriminate between the pair types at a level comparable to healthy controls, with Identical pairs rated the highest, followed by Similar pairs, and Different pairs rated the lowest.

### 2.4 Data analyses

Data preprocessing was conducted in Python, and statistical analyses were performed in R.

#### 2.4.1 The effect of age and education on SL performance

Ideally, demographic variables that differ significantly between groups should be included as covariates to account for potential confounding effects. In the present study, years of education differed significantly between the TLE and control groups (see **Table 2**) and would therefore ordinarily be considered for inclusion in the statistical models. However, years of education was highly collinear with group membership (TLE versus controls), making it difficult to disentangle the unique contributions of education and group to SL performance. Including both variables in the same model would substantially reduce the ability to estimate their independent effects.

Thus, as an initial step to determine whether education influenced performance on the SL tasks, exploratory Pearson’s correlations were conducted within the healthy control group between education and SL performance measures from each task: RT prediction score, familiarity rating composite score, and overall accuracy on the 2AFC. No significant correlations were observed (all *p*’s > .05), providing evidence that years of education did not influence SL performance within controls. Furthermore, no study to our knowledge has reported contributions of education to SL performance.

Similar correlations were also conducted between age, a demographic variable that was closely matched between the groups (see **Table 2**), and the same SL measures within the control group. Although no significant correlations were observed (all *p*’s > .05), previous work has shown that age can impact SL performance, particularly the explicit expression of SL (Wang, Köhler, et al., 2023). Age has also been associated with progressive volume reduction in the hippocampus (Raz et al., 2004). For consistency across models, age was therefore included as a covariate in all subsequent analyses. Since the goal of this study was not to determine the effects of age on SL performance, age was treated as a covariate of no interest and was included only to account for its potential contribution to SL performance.

#### 2.4.2 Target detection task

For each participant, average RTs were calculated for correct detections (hits) of target syllables in each syllable position (first, second, and third). Consistent with previous studies, responses were considered hits if they occurred within 1200ms of the target syllable onset (Batterink & Paller, 2017, 2019; Herrera-Chaves et al., 2026; Wang, Köhler, et al., 2023; Wang, Rosenbaum, et al., 2023). All other keyboard responses were considered false alarms. Misses were classified as any target syllable without a response made within the 1200ms window. To measure participants’ accuracy in detecting the target syllables, a hit rate was calculated as the total number of hits divided by the total number of targets across the task (Hit rate = hits / 144). An RT prediction score was also calculated as a composite measure of facilitation across syllable positions controlling for baseline RTs (Batterink & Paller, 2019). The RT prediction score was calculated by subtracting each patients’ average RT to third-position syllables from their average RT to first-position syllables, then dividing the difference by the average RT to first-position syllables [RT prediction score = (Pos1_RT_avg_ – Pos3_RT_avg_) / Pos1_RT_avg_].

To compare SL performance between TLE patients and healthy controls, two separate repeated measures analyses of covariance (RM ANCOVAs) on mean RTs and hit rates were conducted with group (TLE, Controls) as a between-subjects factor, syllable position (1^st^, 2^nd^, 3^rd^) as a within-subject factor, and age as a covariate.

As an additional analysis to compare RT facilitation between TLE patients and healthy controls, an independent samples *t*-test was conducted with group (TLE, Controls) as the grouping variable and RT prediction score as the dependent variable.

#### 2.4.3 Familiarity rating task

Average familiarity ratings by word type (Word, Partword, Nonword) were calculated for each participant. To assess sensitivity in discriminating target from foil items, two difference scores were computed for each participant by subtracting the average Partword and Nonword ratings from the average Word rating: (W-PW score = W_avg_ – PW_avg_; W-NW score = W_avg_ – NW_avg_). A familiarity rating composite score was also calculated for each participant as an overall measure of discrimination between Words and foil items (Batterink & Paller, 2019) by subtracting the average of both Partword and Nonword ratings from the average Word rating: (FR score = W_avg_ – (PW_avg_ + NW_avg_)/2).

To compare average familiarity ratings across word types and between TLE patients and healthy controls, a RM ANCOVA was conducted with group (TLE, Controls) as a between-subjects factor, item type (W_avg_, PW_avg_, NW_avg_) as a within-subject factor, and age as a covariate.

#### 2.4.4 Two-alternative forced-choice task

For each participant, accuracy was calculated for each condition type (Words vs. Partwords, Words vs. Nonwords). Accuracy was computed as the proportion of correct responses relative to the total number of trials (Condition Accuracy = correct responses / 16).

To compare accuracy between TLE patients and healthy controls, a RM ANCOVA was conducted with group (TLE, Controls) as a between-subjects factor, the two trial type accuracy scores (W vs. PW accuracy, W vs. NW accuracy) as a within-subjects factor, and age as a covariate. Furthermore, one-sample t-tests were conducted to determine whether both healthy controls’ and TLE patients’ performance significantly differed from chance level (proportion correct = .50) across the task overall and separately for each trial type.

#### 2.4.5 Perceptual similarity task

Average similarity ratings by pair type (Identical, Similar, Different) were calculated for each participant. To assess sensitivity in discriminating between target and foil pairs, two difference scores were computed for each participant by subtracting the average Similar and Different pair ratings from the average Identical pair rating: (I-S score = I_avg_ – S_avg_; I-D score = I_avg_ – D_avg_).

To compare sensitivity in discriminating target pairs from foil pairs in TLE patients and healthy controls, two separate ANCOVAs were conducted with group (TLE, Controls) as a fixed factor, either the I-S or I-D difference score as the dependent variable, and age as a covariate.

## 3 Results

For all analyses, Greenhouse-Geisser corrections were reported for factors with more than two levels whenever Mauchly’s test indicated a violation of sphericity.

### 3.1 Target detection task

The TLE group exhibited slower overall RTs compared to healthy controls (Group: *F*(1, 58) = 74.83, *p* < .001, η^2^ = .49). However, both groups responded faster to later-positioned syllables within words (Syllable Position: *F*(2, 116) = 3.16, *p* = .049, η^2^ = .01; linear contrast for syllable position: *t*(58) = -13.87, *p* < .001) and this RT facilitation effect was comparable between the two groups (Syllable Position x Group: *F*(2, 116) = 1.50, *p* = .228, η^2^ = .00; **Figure 5A**). Consistent with this finding, RT prediction scores did not differ significantly between groups (*t*(59) = 1.84, *p* = .071; **Figure 5B**).

**Figure 5:**
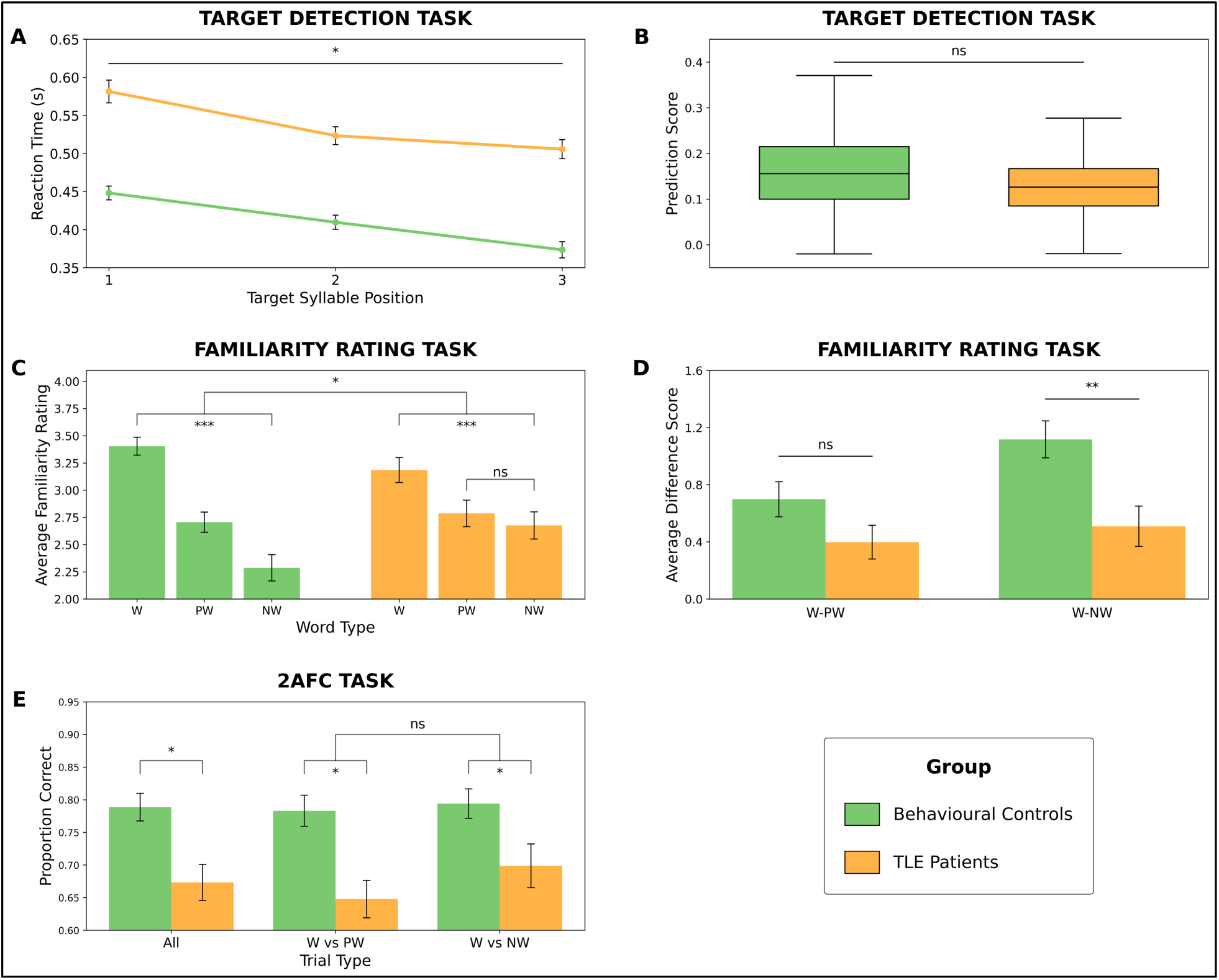
Behavioural results on the statistical learning tasks. (A) Average reaction time in seconds by syllable position on the target detection task. (B) Average reaction time prediction scores by group on the target detection task. (C) Average familiarity ratings by word type on the familiarity rating task. (D) Average difference between familiarity ratings for Words and Partwords (W-PW) and between Words and Nonwords (W-NW) on the familiarity rating task. (E) Average proportion of correct responses on the two-alternative forced-choice (2AFC) task across all trials (All), on trials where Words were paired against Partwords (W vs PW), and on trials where Words were paired against Nonwords (W vs NW). Error bars represent the standard error of the mean. ns = *p* > .05, * = *p* < .05, ** = *p* < .01, *** = *p* < .001

The TLE patient group also exhibited a significantly lower detection accuracy (*M* = 69.3%) than healthy controls (*M* = 85.5%; Group: *F*(1, 58) = 29.56, *p* < .001, η^2^ = .24). However, detection accuracy did not vary across target syllable positions (Syllable Position: *F*(2, 116) = 1.22, *p* = .296, η^2^ = .00) and this pattern was consistent across groups (Syllable Position x Group: *F*(2, 116) = 2.30, *p* = .109, η^2^ = .01).

Overall, these findings suggest that, despite reduced detection accuracy and slower processing speed, TLE patients showed preserved sensitivity to the statistical structure of the stimuli, as evidenced by prediction of upcoming syllables.

### 3.2 Familiarity rating task

Across both groups, Words were rated as the most familiar, followed by Partwords, and Nonwords rated as the least familiar (Word Type: *F*(2, 116) = 10.92, *p* < .001, η^2^ = .07). However, the two groups differed significantly in their pattern of familiarity ratings across word types (Group x Word Type: *F*(2, 116) = 4.73, *p* = .011, η^2^ = .03). Within-group comparisons revealed that controls rated Words as more familiar than both Partwords (*t*(58) = 5.94, *p* < .001) and Nonwords (*t*(58) = 8.70, *p* < .001), and Partwords as more familiar than Nonwords (*t*(58) = 3.48, *p* = .003). Similarly, TLE patients rated Words as more familiar than both Partwords (*t*(58) = 3.06, *p* = .009) and Nonwords (*t*(58) = 3.82, *p* < .001), but the difference in ratings between Partwords and Nonwords was not significant (*t*(58) = 1.07, *p* = .536; **Figure 5C)**.

To probe this interaction further, we conducted separate ANCOVAs on W-PW and W-NW difference scores to test for group differences in discriminating Words from the two foil items. A significant difference was observed for W-NW scores (Group: *F*(1, 58) = 8.49, *p* = .005, η^2^ = .12), with TLE patients showing a smaller discrepancy between ratings for Words and Nonwords compared to healthy controls. In contrast, W-PW scores showed no significant group differences (Group: *F*(1, 58) = 2.75, *p* = .103, η^2^ = .05; **Figure 5D).**

Overall, these findings suggest broadly that TLE patients were able to discriminate between Words and foil items (Partwords and Nonwords), but not to the same extent as healthy controls.

### 3.3 Two-alternative forced-choice task

TLE patients showed poorer accuracy than healthy controls overall (Group: *F*(1, 42) = 5.35, *p* = .026, η^2^ = .09) and this group difference was consistent across trial types (Trial Type: *F*(1, 42) = .00, *p* = .954, η^2^ = .00; Trial Type x Group: *F*(1, 42) = .73, *p* = .399, η^2^ = .00; **Figure 5E**).

However, both groups performed above chance overall (Controls: *t*(34) = 13.6, *p* < .001; TLE: *t*(10) = 4.01, *p* = .002) and on both trial types, in which Words were paired against either Partwords (Controls: *t*(34) = 11.9, *p* < .001; TLE: *t*(10) = 3.30, *p* = .008) or Nonwords (Controls: *t*(34) = 13.1, *p* < .001; TLE: *t*(10) = 3.79, *p* = .004).

Overall, these findings suggest that, although TLE patients demonstrated some learning on this task, they exhibited a relative impairment overall in forced-choice recognition.

### 3.4 Perceptual similarity task

Both groups showed the expected pattern of ratings, with Identical sound pairs rated as the most similar (TLE: *M* = 3.90, Controls: *M* = 3.93), followed by Similar sound pairs (TLE: *M* = 2.20, Controls: *M* = 2.35), and Different sound pairs rated as the least similar (TLE: *M* = 1.04, Controls: *M* = 1.07). Difference score analyses revealed no significant group differences in discriminating between Identical and Similar pairs (Group: *F*(1, 40) = .36, *p* = .553, η^2^ = .01) nor between Identical and Different pairs (Group: *F*(1, 40) = .001, *p* = .980, η^2^ = .00).

Overall, these findings provide behavioural evidence that suggest TLE patients and controls can perceive and discriminate between similar-sounding auditory stimuli to a comparable degree.

## 4 Discussion

The present study investigated whether implicit and explicit expressions of SL are dissociable in TLE patients. Consistent with this hypothesis, TLE patients showed preserved implicit SL expression, exhibiting comparable RT facilitation across syllable positions during the target detection task, alongside impaired explicit expressions of SL. Specifically, they showed an overall deficit in discriminating between Words and both foil types on the 2AFC task, and in particular showed reduced strength of discrimination between Nonwords and Words on the familiarity rating task. Together, these findings suggest that TLE selectively impairs explicit memory, but not implicit memory, of regularities acquired through SL.

The behavioural findings indicate that TLE patients had difficulties intentionally retrieving learned regularities across both explicit measures of SL. On the familiarity rating task, TLE patients did not show a significant difference in their ratings of Partwords and Nonwords. Further examination revealed reduced differentiation specifically for Nonwords relative to Words, with TLE patients erroneously assigning them higher familiarity ratings than healthy controls. This resulted in a higher baseline for unfamiliar foil items in the TLE group. One possibility is that TLE patients relied more heavily on familiarity for the individual syllables encountered in the speech stream, rather than the higher-order statistical structure, when evaluating foil items. Since both Partwords and Nonwords were composed of syllables that appeared during the exposure phase but were recombined into partially intact or fully non-intact sequences, reliance on lower-level familiarity cues may have reduced their sensitivity to the structural differences between the two foil types. Similarly, on the 2AFC task, TLE patients showed reduced accuracy overall in discriminating Words from foil items compared to healthy controls, regardless of whether it was paired against a Partword or Nonword. However, the smaller sample size for the 2AFC task compared to the other SL tasks limits our ability to strongly interpret these findings. Importantly, however, learning was still evident in the TLE group on both explicit measures. They rated Words as being more familiar than both foil types on the familiarity rating task and demonstrated above-chance recognition of Words relative to foils on the 2AFC task. In sum, while the TLE group still showed above-chance evidence of learning on our explicit measures, they were relatively impaired in their explicit expression of SL compared to the control group.

In contrast, the implicit expression of SL was preserved in TLE patients relative to controls. Both groups exhibited comparable RT facilitation across syllable positions during the target detection task, and this was further supported by no significant differences between groups on their prediction scores. These findings suggest that patients with TLE were able to extract the regularities from the continuous speech stream and use that knowledge to predict later positioned syllables within words during real-time processing, despite showing deficits on the explicit measures of SL. Although TLE patients responded more slowly overall and exhibited poorer detection accuracy compared to controls, these differences are unlikely to reflect an impairment in the implicit expression of SL. Rather, they are consistent with generalized cognitive and motor slowing that is commonly reported in TLE (Hwang et al., 2019) and other clinical populations (e.g., Arroyo et al., 2021; Denney et al., 2004; Mathias & Wheaton, 2007). Critically, group differences were absent for the primary learning measure – the RT facilitation across syllable positions – indicating preserved implicit expression of statistical knowledge.

The observed dissociation between preserved implicit and impaired explicit expressions of SL in TLE patients has important implications for understanding the neural mechanisms supporting these forms of memory retrieval. Schapiro et al.’s (2017) computational model proposed that SL depends on the monosynaptic pathway extending from the entorhinal cortex to CA1. However, this model does not differentiate hippocampal contributions to the acquisition versus expression of statistical knowledge, nor does it account for differences in how statistical knowledge is probed. It therefore assumes that this pathway is necessary for all aspects of SL. In partial support of this model, we observed deficits in the TLE group only on the explicit measures of SL, while their performance on an implicit measure of SL remained intact. Our findings thus lend support for an additional dimension to this model, whereby the hippocampus may only be necessary for the explicit expression of SL, but not the learning or implicit expression of SL. This interpretation is consistent with previous lesion-based studies demonstrating impairments on explicit measures of SL following hippocampal damage (Aljishi et al., 2024; Covington et al., 2018; Schapiro et al., 2014; Wang, Rosenbaum, et al., 2023), alongside preserved performance on an implicit measure of SL in one case (Wang, Rosenbaum, et al., 2023). Similar dissociations between implicit and explicit memory processes have also been reported more broadly in TLE (Billingsley et al., 2002; Del Vecchio et al., 2004).

The role of CA1 in SL is particularly interesting because it serves as the convergence point for both the monosynaptic and trisynaptic pathways. CA1 is thought to act as a key integrative hub within the hippocampus, receiving convergent inputs from both the trisynaptic pathway via CA3 and directly from the entorhinal cortex through the monosynaptic pathway. Through this convergence, CA1 may integrate newly encountered information with existing memory representations, supporting the detection of similarities and discrepancies between them (Duncan et al., 2012; Schlichting et al., 2014). However, it remains unclear whether other regions that are part of the trisynaptic pathway, including the dentate gyrus and CA3, are involved in SL. A previous study of SL in a patient with a highly selective bilateral lesion to the dentate gyrus demonstrated impairments in both pattern separation (the proposed function of the trisynaptic pathway; Schapiro et al., 2017) and explicit expression of SL, suggesting that dentate gyrus integrity may also contribute to performance on explicit tasks (Wang, Rosenbaum, et al., 2023). Consequently, our findings cannot exclude the role of the trisynaptic pathway in SL. Future work combining measures of hippocampal subfield integrity with behavioural measures of SL and pattern separation could help determine whether these processes rely on shared or dissociable components of the hippocampus.

This pattern of preserved implicit, but impaired explicit, memory in TLE aligns with traditional neuropsychological theories of long-term memory, which identify the medial temporal lobe, including the hippocampus, as critical for explicit memory, whereas implicit memory depends on neural systems outside the medial temporal lobe (Squire & Zola, 1996). Our findings help reconcile these two perspectives – Schapiro et al.’s (2017) proposal that hippocampal contributions are critical for SL and Squire and Zola’s (1996) theory that the hippocampus supports explicit but not implicit memory – by suggesting that the hippocampus contributes selectively to the explicit expression of statistical knowledge, while implicit predictive processing can remain intact despite hippocampal abnormalities. Complementary evidence for this neural dissociation comes from intracranial EEG studies, which often show robust entrainment to statistical regularities in modality-specific cortical regions during exposure, but not within the hippocampus (Henin et al., 2021; Herrera-Chaves et al., 2026). These findings suggest that statistical regularities can be acquired by cortical systems that may subsequently support implicit behavioural expression without requiring hippocampal involvement.

Although the present findings align with existing accounts that implicate hippocampal abnormalities in behavioural expression of statistical knowledge, they should not be interpreted as demonstrating that the hippocampus acts in isolation. TLE is increasingly recognized as a network disorder involving structural and functional abnormalities extending beyond the hippocampus to encompass broader medial and lateral temporal regions and their connections with distributed cortical networks (Bernhardt et al., 2013). Previous neuroimaging studies using fMRI and intracranial EEG have also implicated broader cortical and subcortical structures in SL, including the insula, striatum, the inferior frontal gyrus, the middle and superior temporal gyri, the supramarginal gyrus, and the occipital cortex (Herrera-Chaves et al., 2026; Karuza et al., 2013; McNealy et al., 2006; Sandoval et al., 2017; Schapiro et al., 2012; Turk-Browne et al., 2009). This raises the possibility that broader abnormalities outside the hippocampus or even beyond the temporal lobe also contributed to the impairments observed in explicit SL expression in our study. Importantly, TLE patients performed comparably to controls on the perceptual similarity task, indicating that the observed deficits in the explicit expression of SL cannot simply be attributed to lower-level processes of impaired auditory perception or basic sound discrimination. Nevertheless, because the present study did not directly examine the neural structures or networks that underlie this behavioural dissociation, future work incorporating neuroimaging methods will be necessary to determine whether the results observed in this study are directly related to hippocampal integrity, and how the hippocampus interacts with broader networks to support different expressions of SL.

Hemispheric lateralization of language processing and verbal memory may also be particularly relevant given that the present study employed an auditory linguistic SL paradigm. Previous research has demonstrated that patients with TLE often show left hemispheric dominance for language, consistent with neurotypical populations, although they can exhibit more variable profiles depending on the laterality of seizure onset, the age at which seizures began, and handedness (Milner et al., 1964; Neudorf et al., 2020; Springer et al., 1999). Furthermore, left temporal lobe pathology in TLE has been associated with greater memory impairments for verbal material compared with right temporal pathology (Savage et al., 2002; Zaidel et al., 1994; Zaidel et al., 1998). Unfortunately, the small sample sizes of our TLE subgroups limited our ability to directly evaluate whether seizure onset laterality relates to SL performance. Future work should consider these factors alongside neuroimaging measures when examining the neural basis of SL expression for linguistic material.

Other limitations of our study should also be acknowledged. First, our TLE and control groups differed significantly in their levels of education. Although education was not associated with SL performance in our control sample, and no other study that we are aware of has identified education as a predictor of SL ability, future studies should seek to match their participant groups more closely on this variable. Additionally, testing environments differed between these two groups. Patients were assessed within a busy hospital setting where environmental distractions were difficult to control, whereas control participants completed testing in a quiet laboratory environment with minimal distractions. However, although testing conditions were not ideal, our efforts to reduce the impact of environmental noise on patient performance using in-ear headphones appeared to be reasonably effective, as patients performed comparably to controls on the sound discrimination task and showed evidence of learning on the implicit SL task. Nevertheless, future studies should aim to standardize testing environments across groups where possible.

In conclusion, the present study provides behavioural evidence that implicit and explicit expressions of SL are dissociable in TLE patients. Patients with TLE exhibited preserved implicit predictive processing despite also showing impairments in explicit retrieval of learned regularities. These findings refine current computational accounts of the role of the hippocampus in SL (Schapiro et al., 2017) and bridge them together with neuropsychological theories of long-term memory (Squire & Zola, 1996) by suggesting that hippocampal integrity may be particularly important for the explicit expression of statistical knowledge, while implicit expression of SL can remain relatively preserved.

